# Menstrual cycle phase and premenstrual symptoms in relation with the stress response: an fMRI study

**DOI:** 10.64898/2026.09.21.753084

**Authors:** Elise Bücklein-Ehlers, Ann-Christin S. Kimmig, Thomas Dresler, Michael Lämmerhofer, Tamara Janker, Lydia Kogler, Birgit Derntl, Andreas J. Fallgatter, Erika Comasco, Ann-Christine Ehlis

## Abstract

**Objective:** Women with premenstrual syndrome (PMS) experience affective lability, irritability, depressed mood, and anxiety during the late luteal phase of the menstrual cycle. Stress reactivity may contribute to the development of these symptoms, yet neural correlates remain unexplored. We examined neural, behavioral and endocrine stress responses across menstrual cycle phases in women with and without PMS.

**Methods:** 26 women with PMS and 24 women without PMS participated in two fMRI sessions, during the mid-follicular (days 5 to 11) and late-luteal phases (days −7 to −1). Participants completed the Montreal Imaging Stress Task (MIST). Subjective stress ratings and salivary cortisol were collected at six time points before and after stress induction. Stress-related brain activation was assessed using whole-brain and region-of-interest analyses.

**Results:** Subjective stress ratings increased as expected from before to after the MIST, with no group or phase differences. Salivary cortisol was significantly higher during the mid-follicular phase, with no group effect or significant pre-post MIST difference. While the stress task elicited activation patterns in stress-related regions, whole-brain analysis revealed no significant effects of group (PMS vs. no PMS) or menstrual cycle phase (follicular vs. luteal) on the neural stress response. Considering premenstrual symptom severity as a predictor revealed significant group-by-symptom severity interactions in the right inferior frontal gyrus and the right superior temporal gyrus.

**Conclusion:** Using a within-subject repeated-measures design, we found neural responses to psychosocial stress to be stable across menstrual cycle phases. Individual differences in symptom severity may nevertheless be associated with differences in stress-related brain activation.

## Introduction

Psychosocial stress is commonly experienced and usually elicits a stress response on a neural, behavioral, as well as endocrine level. Central to the stress response is the brain, where stressors and available resources to cope with them are evaluated and integrated to coordinate an appropriate response (McEwen, 2017). Key regulator of the stress response on an endocrine level is the hypothalamus-pituitary-adrenal (HPA) axis, which, through a cascade of hormonal signaling, leads to the release of cortisol into the bloodstream, from where it can reach all organs (Charmandari et al., 2005).

Meta-analyses consistently identify activations of the bilateral insula, the inferior frontal gyrus (IFG), the superior temporal gyrus (STG) as well as the thalamus during psychosocial stress, together with deactivation of the dorsal striatum (caudate nucleus) (Berretz et al., 2021; Kogler et al., 2015b). Complementing these meta-analytic findings, studies using functional near-infrared spectroscopy (fNIRS) during the Trier Social Stress Test (TSST) have linked stress-related changes in IFG and dorsolateral prefrontal cortex (dlPFC) activity to rumination in healthy females and males (Rosenbaum et al., 2018b, 2018a).

Another endocrine system that interacts with the HPA-axis is the hypothalamic-pituitary-gonadal (HPG) axis, which regulates ovarian hormone fluctuations associated with the menstrual cycle. The menstrual cycle is commonly divided into the follicular phase, characterized by low levels of progesterone and increasing levels of estradiol, and the luteal phase, during which both hormones initially increase before declining prior to menstruation (Schmalenberger et al., 2021).

Menstrual cycle phases are known to influence cortisol levels in naturally cycling women, with two meta-analyses reporting elevated basal cortisol levels during the follicular compared to the luteal phase (Hamidovic et al., 2020; Klusmann et al., 2022), and one meta-analysis reporting slightly increased cortisol reactivity in response to stress during the luteal compared to the follicular phase (Klusmann et al., 2023).

On a neural level, few studies have investigated stress responses across menstrual cycle phases. Chung et al. (2016) compared women in the early follicular and mid-luteal phase using the Montreal Imaging Stress Task (MIST), a validated fMRI paradigm for inducing psychosocial stress (Dedovic et al., 2005), and found higher hippocampal activation during the mid-luteal phase compared with the early-follicular phase. Similarly, Albert et al. (2015) investigated the impact of endogenous estradiol on neural responses to psychosocial stress using the MIST, and observed reduced deactivation in limbic regions during ovulation compared with the early follicular phase. Furthermore, Ossewaarde and colleagues (2010) reported menstrual cycle-related effects on stress sensitivity in the amygdala during the late luteal compared with the late follicular phase, using aversive movie clips for inducing stress. Although these studies provide initial evidence for menstrual cycle influences on neural stress processing, two of them are limited by their between-subject design, leaving within-person changes across the menstrual cycle largely unexplored.

Up to 90% of individuals with a menstrual cycle report cyclical physical or affective symptoms, but these are typically mild and do not interfere with daily functioning (Braverman, 2007). In contrast, premenstrual disorders (PMDs), comprising of premenstrual syndrome (PMS) and the more severe Premenstrual Dysphoric Disorder (PMDD), are characterized by recurrent symptoms that impair daily functioning (Halbreich et al., 2003). PMS affects around 15-30% of the menstruating population (Halbreich et al., 2003; Yonkers and Simoni, 2018), whereas PMDD affects 3-8% (Epperson et al., 2012; Reilly et al., 2024; Yonkers and Simoni, 2018). Affective symptoms include irritability or anger, mood lability, anxiety, and depressive symptoms, but physical symptoms, such as breast swelling, bloating, or fatigue, are also common (Halbreich et al., 1982). These symptoms emerge during the luteal phase, and remit shortly after the onset of menstruation (Epperson et al., 2012). Current research suggests that not altered levels, but instead hypersensitivity to menstrual cycle-related fluctuations in ovarian hormones may underlie these symptoms (Comasco et al., 2021; Rubinow and Schmidt, 2025; Schmidt et al., 2017). Although the precise etiology of PMDs remains unclear, dysregulation of the HPA axis has emerged as a potential contributing mechanism. Several studies have reported attenuated HPA axis function in PMDs, though not all studies have consistently observed these effects, potentially reflecting methodological differences across studies (Hantsoo et al., 2023; Kiesner and Granger, 2016).

Evidence of altered HPA axis functioning show blunted cortisol responses to stress in women with PMDs (Hamidovic et al., 2024; Huang et al., 2015), flatter diurnal cortisol slopes and delayed cortisol awakening response (CAR) peaks (Beddig et al., 2019), as well as an overall attenuated CAR (Hoffmann et al., 2025; Hou et al., 2019). A recent meta-analysis further demonstrated that women with PMDs report higher levels of perceived stress, particularly during the luteal phase, and that a history of trauma is linked with an approximately two-fold increased likelihood of experiencing PMDs (Bencker et al., 2025).

Despite accumulating evidence linking PMDs with alterations in stress-related endocrine functioning, the neural correlates of the stress response in relation to both, menstrual cycle phase and premenstrual symptoms remain unknown.

The present study therefore aimed to investigate the response to psychosocial stress in women with and without PMS using a repeated-measures design in which participants were assessed during both the mid-follicular and late-luteal phase. We examined the stress response from neural, behavioral and endocrine perspectives, operationalized as task-based brain activity, subjective stress ratings, and salivary cortisol concentrations, respectively.

We expected psychosocial stress to elicit activation effects in key regions of the stress network, including the bilateral insula, thalamus, IFG, STG, and deactivation effects in the parahippocampal gyrus, the dlPFC as well as the left caudate nucleus (Berretz et al., 2021; Kogler et al., 2015b; Rosenbaum et al., 2018a, 2018b). We further hypothesized that stress-related neural responses would vary across menstrual cycle phases, due to modulatory effects of ovarian hormones on stress responses (Albert et al., 2015; Chung et al., 2016; Ossewaarde et al., 2010), and between women with and without PMS, although the direction of PMS-related effects remained exploratory.

## Material and Methods

### Participants

Participants were recruited through e-mail to university students and staff, as well as flyers around town and in gynecological doctors’ offices. Participants first attended a screening visit in which they received information on the study and gave written consent. Participants were reimbursed for their participation in the study with 100€. All participants were German speaking and had to have regular menstrual cycles (between 25 and 35 days; no signs of menopause). They were excluded if they had any current neurological or mental disorder, confirmed via standardized clinical interview for DSM-5, clinical version SCID-5-CV (Beesdo-Baum et al., 2019). They were furthermore excluded if they used psychotropic medication, were on steroid treatment or hormonal contraception during the previous three months, had severe medical conditions (such as hormonal, metabolic or chronic diseases), were pregnant or breastfeeding in the last 12 months, smoked regularly (more than 3 cigarettes per week), or had any contraindications for MRI. Subjects participating in competitive sports or shift work were also excluded due to the effects on cortisol and stress reactivity (King and Liberzon, 2009).

### PMS grouping procedure

Group assignment was based on a retrospective assessment of premenstrual symptoms using a self-report questionnaire describing criteria for PMDD from the DSM-5 (American Psychiatric Association, 2013). For the full questionnaire, please refer to the Supplementary Material (S1). All participants subsequently completed prospective daily ratings of premenstrual symptoms for one to two menstrual cycles, depending on their group assignment, using the Daily Record of Severity of Problems (DRSP) (Endicott et al., 2006). Questionnaires were used for a prospective assessment of premenstrual symptom severity. The DRSP consists of 21 items, assessing the 11 affective and somatic symptoms of PMDD according to DSM-5 criteria. Each item is rated on a scale from 1 (“not at all”) to 6 (“extreme”). Participants completed the questionnaire via the m-path application (Mestdagh et al., 2023) on their personal smartphones, with daily reminders sent at 8 pm.

### MRI Procedure

All participants were scheduled for two MRI measurements, once in the mid-follicular phase (day 4 to 10 after beginning of menses) and once in the late luteal phase (days −7 to −1 before suspected beginning of the next menses). An overview of the study design is depicted in Figure 1A. Late luteal phase testing was confirmed using self-reported first days of the current and subsequent menstrual cycle as well as ovulation testing via LH test strips (nal von minden GmbH, Moers, Germany; sensitivity 30 mlU/ml hLH) between days −12 and −16, counting back from the expected onset of next menses (Schmalenberger et al., 2021). Measurement order was randomized to avoid order effects: 50% of controls and 54% of PMS participants underwent their first scan in the mid-follicular phase. Measurements always took place on workdays and started in the afternoon between 2 and 6 pm to ensure similar levels of cortisol. To control for confounds in salivary cortisol, participants were asked to avoid caffeine and exercise for 3 hours, food, sugary drinks and chewing gum for 2 hours, and smoking, alcohol, and medication for 24 hours before testing.

**Figure 1.**
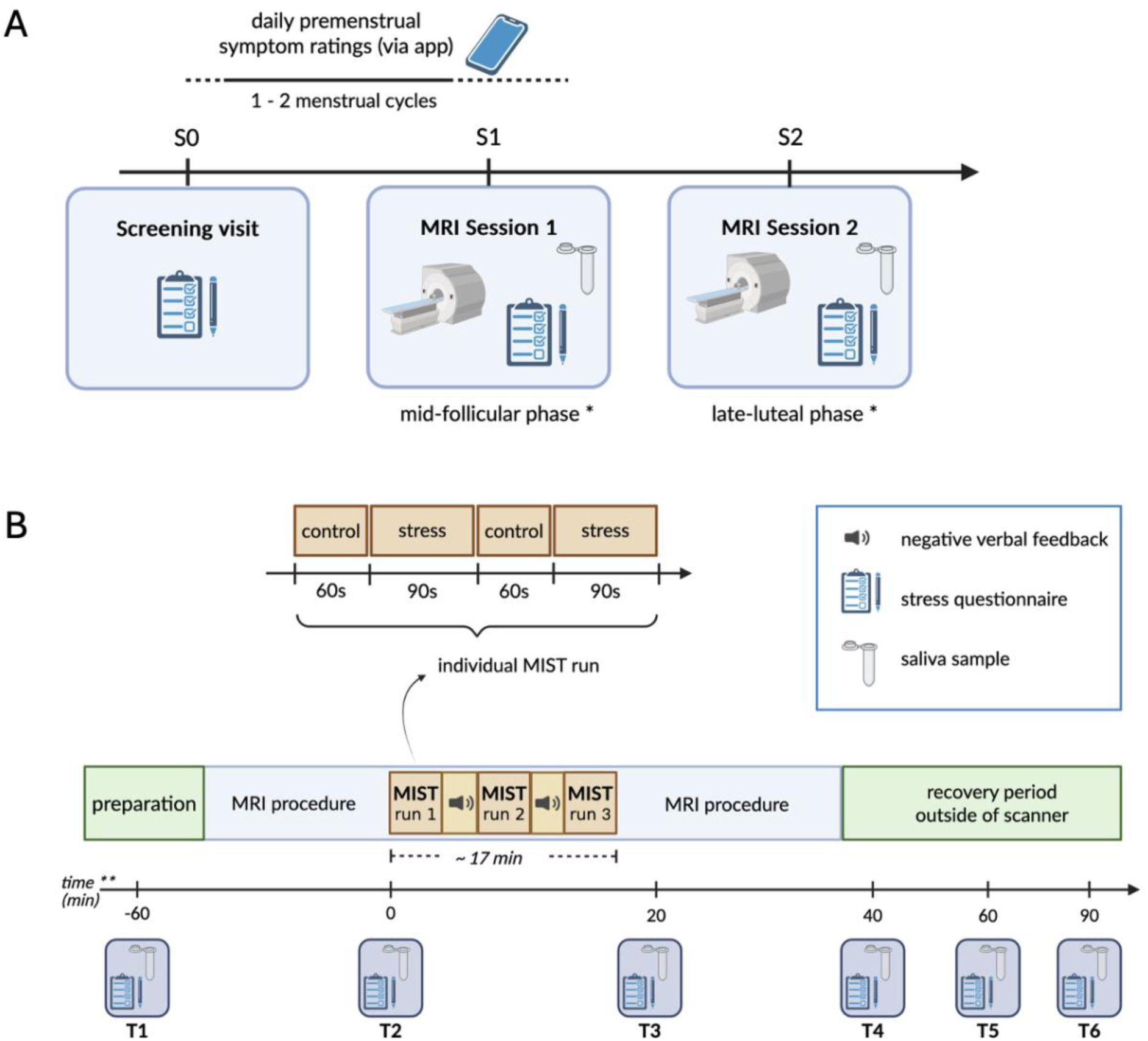
Study procedure including illustration of MRI sessions with the Montreal Imaging Stress Task. ***Notes.*** Figure A shows the study procedure and Figure B the MRI procedure on measurement days, with special focus on the Montreal Imaging Stress Task procedure. ** time arrow not in scale. MIST = Montreal Imaging Stress Task; MRI = Magnetic resonance imaging.

The study was approved by the ethical committee of the Medical Faculty of University of Tübingen (approval number: 340/2023BO2), preregistered at https://clinicaltrials.gov (trial no. NCT06130371) and carried out in accordance with the ethical standards of the Helsinki Declaration.

### Experimental stress paradigm: Montreal Imaging Stress Task

Psychosocial stress was induced using the Montreal Imaging Stress Task (MIST) (Dedovic et al., 2005), in which participants solve arithmetic problems presented on a computer screen by selecting one-digit numbers from a rotary dial while undergoing fMRI. During the experimental condition, a time-limit progress bar and a performance bar showing participant’s scores relative to a (pre-set) average are displayed to induce stress. The difficulty of the task is adapted in a way that the subject reaches no more than 45-50% correct answers. During the control condition, the arithmetic tasks are presented without time restraints and without negative feedback (Dedovic et al., 2005).

To familiarize the participants with the task, the MIST was practiced shortly before the MRI session on a laptop for 90 seconds with the control condition. During the MRI session, the MIST was presented in a block design with three runs. Each run consisted of two repetitions of control and experimental blocks for 60 and 90 seconds, respectively. In between the runs, the investigator provided negative feedback via the intercom system, urging the participant to perform better to increase psychosocial stress. For an illustration of the MRI procedure as well as the MIST, see Figure 1B.

### General self-report questionnaires

Socio-demographic data as well as questionnaire data regarding trait anxiety (State-Trait Anxiety Inventory – Trait version, STAI-T, Spielberger, 1983) and childhood trauma (Childhood Trauma Questionnaire, CTQ, Wingenfeld et al., 2010) was acquired during the screening visit, and questionnaire data regarding chronic stress (Trier Inventory of the Assessment of Chronic Stress, TICS, Schulz et al., 2004), perceived stress (Perceived Stress Scale – 10 item version, PSS-10, Cohen and Williamson, 1988; Reis et al., 2019) and depressive symptoms (Beck Depression Inventory-II, BDI-II, Kühner et al., 2007) was collected at both MRI sessions to control for potential confounding effects on stress.

### Subjective stress and salivary cortisol

Throughout the experimental session, participants were asked to rate their affective level on a 10-point Likert scale from 1: not at all to 10: extremely, with five items regarding how (1) stressed, (2) tense, (3) challenged, (4) burdened, (5) struggling they felt. The mean across these five questions was used as the mean subjective stress rating. At the same time as the subjective rating, a saliva sample was collected for cortisol analysis, using a synthetic fiber swab (Salivette® Cortisol, Sarstedt, Nümbrecht, Germany). Subjective stress ratings and salivary cortisol samples were collected at six timepoints: 60 min before the MIST (T1, baseline), directly before (T2), directly after (T3), and 40 (T4), 60 (T5) and 90 min (T6) after the MIST.

Salivary cortisol samples were analyzed at the Institute of Pharmaceutical Sciences, University of Tübingen, using a targeted liquid chromatography-mass spectrometry assay. For a detailed description of the saliva analysis procedure please refer to the supplementary material S2.

### Image acquisition

Functional and anatomical imaging data were acquired at the University Hospital Tübingen with a 3T Siemens MAGNETOM Prisma_XR scanner using a 64-channel head coil, with the forehead fixated using tape to minimize head motion (Krause et al., 2019). First, a T1-weighted high-resolution anatomical image was acquired using an MPRAGE sequence (208 sagittal slices, TR = 2400 ms, TE = 2.22 ms, TI = 1000 ms, voxel size = 0.8 x 0.8 x 0.8 mm, flip angle = 8°, distance factor = 50%, GRAPPA acceleration factor = 2, FOV = 256 mm). Additionally, a T2-weighted SPACE image followed (208 sagittal slices, TR = 3200 ms, TE = 562 ms, voxel size = 0.8×0.8×0.8 mm³, FOV = 256 mm). To account for magnetic field distortions in echo-planar images (EPIs), a T2-weighted B0 inhomogeneity field map was acquired (68 slices, TR = 745 ms, TE = 5.19/7.65 ms, voxel size = 2.3×2.3×2.0 mm³, FOV = 220 mm, flip angle = 60°). During the task, functional images were acquired using a multiband T2-weighted EPI sequence (68 interleaved slices, multiband factor = 4, TR = 1400 ms, TE = 30 ms, voxel size = 2 x 2 x 2 mm, FOV = 220 mm, flip angle = 65°).

### Statistical analyses

Data analysis was run using Statistical Parametric Mapping software (SPM12) (Ashburner et al., 2014) or R (R Core Team, 2023), depending on the research question. If not otherwise specified, two-tailed testing with an α-level of 0.05 was applied.

### Subjective stress and salivary cortisol

To check whether stress was induced on a subjective and endocrine level during application of the MIST, subjective stress and salivary cortisol were analyzed in linear mixed models with all six timepoints (T1-T6, see Fig. 1), menstrual cycle phase (mid-follicular vs. late-luteal) and group (no PMS vs. PMS), using the lme4 package in R (Bates et al., 2015).

### fMRI data preprocessing and analysis

fMRI scans were first preprocessed using the fMRIPrep pipeline (fMRIPrep 24.1.1 (Esteban et al., 2020, 2019; Markiewicz et al., 2024); RRID:SCR_016216). Preprocessing included realignment, co-registration to T1-weighted anatomical image, and spatial normalization to Montreal Neurological Institute (MNI) space. Information on preprocessing with fMRIPrep is provided in full in the supplementary material (S2). In a next step, spatial smoothing was applied using the Statistical Parametric Mapping software (SPM12) (Ashburner et al., 2014), using a 6mm Gaussian kernel (full width at half maximum).

At the first level, a general linear model was estimated for each participant using SPM12 implemented in MATLAB (version R2024a). Design matrices were convolved with the default gamma hemodynamic response function (HRF) and its temporal derivative to account for small variations in response latency. The first six motion parameters were added to the model as covariates of no interest. One participant showed excessive head movement (> 2mm or 2°), but because results of the full factorial design did not change when excluding this participant, and motion parameters were added, all data was kept for the analysis. A high-pass filter (cutoff 256 seconds, due to long blocks of 60-90 seconds) was applied to remove low-frequency drifts from the time series. To investigate the effects of psychosocial stress on brain activation, the control condition was subtracted from the experimental condition and t-contrast maps were generated.

### Whole-brain analysis of group and phase differences in stress-induced brain activity

Second-level analyses were conducted using a 2 x 2 full-factorial design with factors cycle phase (mid-follicular vs. late-luteal) and group (no PMS vs. PMS) on the contrast images. The main effects as well as the interaction effect of group x phase were considered with cluster-level FWE-correction at *p* < 0.05. All emerging clusters were checked with the Anatomy toolbox in SPM12 (Eickhoff et al., 2005).

### ROI analysis of group and phase differences in stress-induced brain activity

A priori ROIs were derived from two meta-analyses on brain activity associated with acute psychosocial stress exposure (Berretz et al., 2021; Kogler et al., 2015b). These meta-analysis-derived ROIs included the left insula (x = −32, y = 20, z= 4), right insula (x = 34, y = 24, z = 2), and thalamus (x = 14, y = 0, z = −2), as well as the right inferior frontal gyrus (IFG; x = 38, y = 22, z = 8) and right superior temporal gyrus (STG; x = 62, y = −40, z = 22) for activation effects. For deactivation effects, we analyzed the parahippocampal gyrus (x = 18, y = −6, z = −18) and left dorsal striatum (caudate nucleus) (x = −10, y = 18, z = −6). In addition, the bilateral dorsolateral prefrontal cortex (dlPFC) (BA 46) was analyzed for deactivation effects, since it has also been implicated in stress processing (Int-Veen et al., 2023; Rosenbaum et al., 2018a).

Meta-analysis-derived ROIs were defined as 8-mm-radius spheres using MarsBaR (Brett et al., 2002). The dlPFC ROI was defined anatomically using the Talairach Daemon atlas (Lancaster et al., 2000) within the WFU Pickatlas toolbox (Maldjian et al., 2003) in SPM12.

ROI data were then extracted and analyzed in MarsBaR (Brett et al., 2002), using the factorial design from SPM12 from the whole-brain analysis.

### Associations between stress-induced brain activity and PMS symptom severity

To examine associations between brain activity and symptom severity, mean luteal DRSP scores were included in the previous models. Scores were averaged across days −7 to −1 to reflect late-luteal symptom severity. DRSP data were collected during screening and did not overlap with MRI measurements.

On the whole-brain level, a full factorial model with the mean DRSP score as an added predictor was calculated with SPM12. For the ROI analysis, linear mixed models were calculated in R with the *lme4* package (Bates et al., 2015) to adjust for multiple measurements per subject. Dependent variables were the selected ROIs, and independent predictors were group (PMS vs. no PMS), cycle phase (follicular vs. luteal), DRSP score as well as the interaction between group and DRSP score.

## Results

### Participant characteristics

We collected repeated-measures data from *n* = 26 participants with PMS and *n* = 24 participants without PMS. Demographic and psychometric characteristics as well as DRSP scores and p-values for group differences are presented in Table 1. No group difference in age, body mass index (BMI), parity, average cycle length, age at menarche, perceived stress (PSS-10), or DRSP scores during the follicular phase emerged (all *p* < 0.05). Participants in the PMS group more often had university degrees (*p* = 0.024) and a history of depression (*p* = 0.005), reported higher trait anxiety (*p* = 0.002) and childhood maltreatment (*p* = 0.008) at screening, and higher chronic stress during both, luteal (*p* = 0.001) and follicular phase (*p* = 0.003). As expected, participants in the PMS group also had higher DRSP scores during the luteal phase (*p* < 0.001).

**Table 1.** Participants characteristics.

| Characteristic | Mean (SD); n (%) |  | p-value |
| --- | --- | --- | --- |
|  | non-PMS *n* = 24 | PMS *n* = 26 |  |
| **Age (years)** | 27.4 (5.7) | 28.0 (5.7) | 0.7<sup>2</sup> |
| **BMI** |  |  |  |
| luteal | 24.4 (9.0) | 23.0 (3.4) | 0.5<sup>2</sup> |
| follicular | 22.8 (3.2) | 22.7 (3.4) | >0.9<sup>2</sup> |
| **Education level** |  |  | 0.024<sup>3*</sup> |
| below university-level | 16 (67%) | 8 (31%) |  |
| with university degree | 8 (33%) | 18 (69%) |  |
| **Nulliparous** | 22 (92%) | 21 (81%) | 0.5<sup>3</sup> |
| **Average cycle length** | 28.9 (2.3) | 28.7 (2.1) | 0.8<sup>2</sup> |
| **Age at menarche (years)** | 12.9 (1.2) | 12.8 (1.5) | 0.8<sup>2</sup> |
| **Mental history** |  |  |  |
| Depressive disorder | 0 (0%) | 9 (35%) | 0.005<sup>3*</sup> |
| Anxiety disorder | 0 (0%) | 2 (7.7%) | 0.5<sup>3</sup> |
| **STAI-T** | 16.4 (3.7) | 20.3 (4.6) | 0.002<sup>2*</sup> |
| **CTQ** | 32 (5) | 39 (12) | 0.008<sup>2*</sup> |
| **TICS** |  |  |  |
| luteal | 64 (31) | 93 (28) | 0.001<sup>2*</sup> |
| follicular | 64 (27) | 89 (28) | 0.003<sup>2*</sup> |
| **PSS-10** |  |  |  |
| luteal | 21.5 (4.9) | 21.9 (4.7) | 0.8<sup>2</sup> |
| follicular | 19.5 (4.7) | 21.3 (6.0) | 0.2<sup>2</sup> |
| **BDI-II** |  |  |  |
| luteal | 6 (7) | 11 (9) | 0.023<sup>2*</sup> |
| follicular | 4 (4) | 11 (8) | <0.001<sup>2*</sup> |
| **DRSP<sup>1</sup>** |  |  |  |
| follicular | 34 (17) | 35 (9) | 0.9<sup>2</sup> |
| luteal | 34 (11) | 46 (14) | <0.001<sup>2*</sup> |
| **Cycle day MRI** |  |  |  |
| follicular | 8.5 (2.0) | 7.2 (1.7) | 0.012<sup>2*</sup> |
| luteal | -5.0 (1.9) | -5.9 (3.4) | 0.3<sup>2</sup> |

<sup>1</sup>based on prospective ratings during one to two menstrual cycles previous to MRI measurements; <sup>2</sup>Welch Two Sample t-test; <sup>3</sup>Pearson's Chi-squared test; \* significant difference between groups, *p* < 0.05.
*Note.* BDI-II = Beck Depression Inventory II; BMI = body mass index; CTQ = Childhood Trauma Questionnaire; DRSP = Daily Rating of Severity of Problems; MRI = magnetic resonance imaging; PMS = premenstrual syndrome; PSS-10 = Perceived Stress Scale (10-item version); SD = standard deviation; STAI-T = state-trait-anxiety inventory (trait version); TICS = Trier Inventory of Chronic Stress.

### Subjective Stress

Mean subjective stress ratings and individual stress curves from six timepoints before and after the MIST are shown in Figure 2. A linear mixed model with group, phase, the interaction between the two and timepoint included as a factor showed that while subjective stress did not differ significantly between group (*ß* = 0.49, *p* = 0.145) or phase (*ß* = −0.13, *p* = 0.700), or their interaction (*ß* = 0.25, *p* = 0.241), timepoints T3 (*ß* = 2.91, *p* < 0.0001), T5 (*ß* = - 0.42, *p* = 0.021) and T6 (*ß* = −0.56, *p* = 0.002) were significant predictors for subjective stress (using T1 as reference). Post-hoc pairwise comparisons revealed that subjective stress was significantly different between the timepoints before and after the MIST (contrast T2-T3, *ß* = - 2.80, *p_corr_* < 0.0001). Therefore, we assume that the MIST did successfully induce subjective stress in our experiment. Detailed results from the linear model and post-hoc pairwise comparisons can be found in the Supplementary Tables S1.1 and S1.2, respectively.

**Figure 2.**
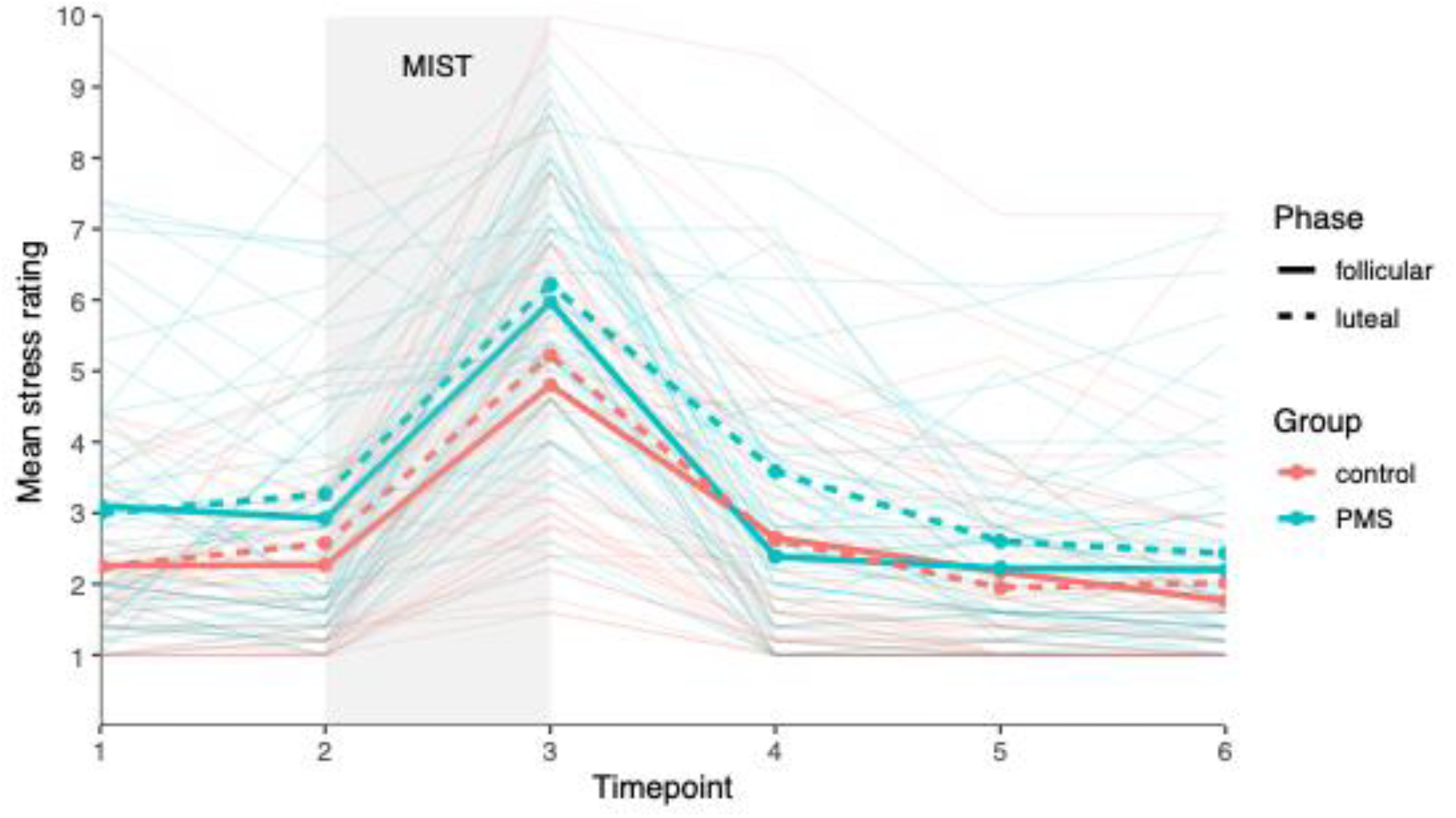
Mean stress ratings across the six timepoints of the MIST. ***Notes.*** Stress was rated on a scale from 1 to 10 at six timepoints: T1 60 minutes before the MIST, T2 directly before the MIST, T3 20 minutes after beginning of the MIST, T4 40 minutes after beginning of the MIST, T5 60 minutes after beginning of the MIST and T6 90 minutes after beginning of the MIST. Individual curves are shown in lighter color in the background. MIST = Montreal Imaging Stress Task, PMS = Premenstrual Syndrome.

### Salivary Cortisol

Descriptive salivary cortisol and individual cortisol curves for the six timepoints are presented in Figure 3. Salivary cortisol was examined for normality and because values were positively skewed, concentrations were log-transformed prior to statistical analyses. A linear mixed model with group, phase, the interaction between the two and timepoint (included as a factor) showed that while the group and the interaction effect were not significant, salivary cortisol was significantly higher in the follicular phase than in the luteal phase (*ß* = −0.17, *p* = 0.027). Timepoint (using T1 as reference) was also a significant predictor, but post-hoc pairwise comparisons showed that salivary cortisol pre-(T2) and post-MIST (T3) did not differ significantly from each other. Detailed results from the linear model and post-hoc pairwise comparisons of all time points can be found in the Supplementary Tables S2.1 and S2.2, respectively.

**Figure 3.**
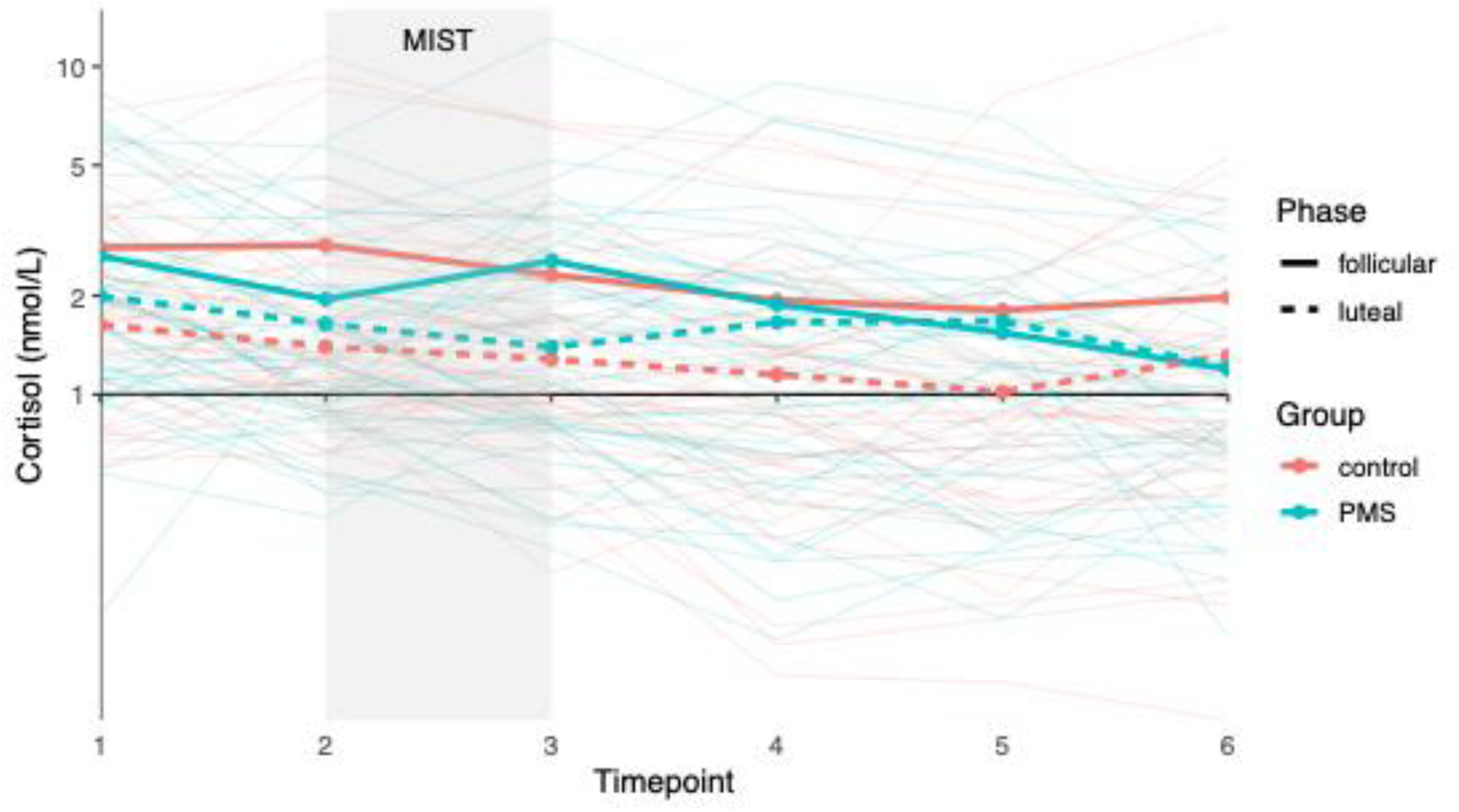
Mean salivary cortisol across the six timepoints of the MIST. ***Notes.*** Salivary cortisol measured at six timepoints: T1 60 minutes before the MIST, T2 directly before the MIST, T3 20 minutes after beginning of the MIST, T4 40 minutes after beginning of the MIST, T5 60 minutes after beginning of the MIST and T6 90 minutes after beginning of the MIST. Individual curves are shown in lighter color in the background. Y-axis displayed on a logarithmic scale to enhance interpretability. MIST = Montreal Imaging Stress Task, PMS = Premenstrual Syndrome.

### Task-based whole-brain fMRI analysis with group and menstrual cycle phase

*Overall effect of stress.* Whole-brain analysis using a full factorial model revealed a significant main effect of stress across all participants and menstrual cycle phases for the contrast stress vs. control (cluster-level FWE-correction at *p* < 0.05). Stress exposure elicited significant activations primarily in a cluster (cluster 1, 58236 vox) spanning the left fusiform gyrus (FuG; x = −24, y = −80, z = −12; *t* = 16.56) and right superior occipital gyrus (SOG; x = 18, y = −96, z = 16; *t* = 15.26), as well as two clusters spanning the bilateral insula (cluster 2, 29881 vox; right: x = 32, y = 20, z = −14; *t* = 7.82; cluster 3, 366 vox; left: x = −32, y = 18, z = −14; *t* = 7.49) and right inferior frontal gyrus (IFG; x = 42, y = 16, z = 4; *t* = 5.80). Significant deactivations were observed in seven clusters. The largest cluster (cluster 1, 7634 vox) included the left IFG (pars triangularis) (x = −52, y = 34, z = 20; *t* = 9.14) and left middle orbital (x = −24, y = 30, z = - 18; *t* = 8.48) and frontal gyrus (x = −52, y = 24, z = 34; *t* = 8.48). Other relevant clusters included the right rolandic operculum (cluster 2, 2822 vox; x = 40, y = −16, z = 18; *t* = 8.91) and right superior temporal gyrus (STG) (x = x = 46, y = −16, z = 0; *t* = 8.17), as well as the left inferior parietal lobule (cluster 3, 1852 vox; x = −34, y = −70, z = 46; *t* = 8.56) and left angular gyrus (x = −38, y = −70, z = 50; *t* = 8.64). Complete cluster statistics, including peak coordinates, cluster sizes and T-values are provided in the Supplementary Tables S3.1 and S3.2 for activation and deactivation clusters, respectively. For an illustration of the effects, see also Figure 4.

**Figure 4.**
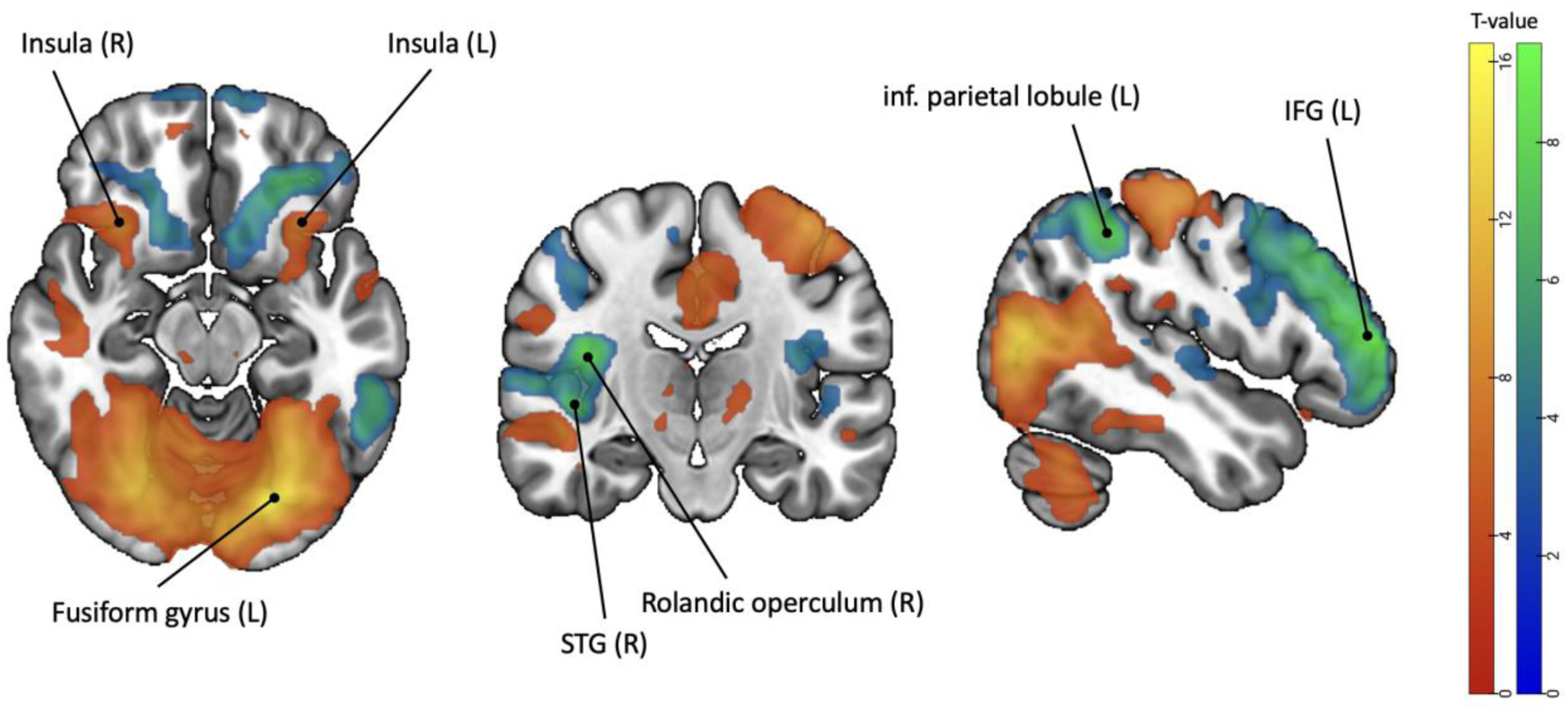
Task-related neural activation and deactivation clusters during the Montreal Imaging Stress Task across all participants, phases and groups. ***Notes.*** Pattern of task-related activations and deactivations from the whole-brain analysis across all subjects, phases and groups for the contrast stress vs. control (*p_FWEc_* < 0.05). IFG = inferior frontal gyrus; inf. = inferior; L = left; R = right; STG = superior temporal gyrus

### No main or interaction effect of group and phase

We found no evidence in our full factorial model of a group (PMS vs. no PMS) or menstrual cycle phase (follicular vs. luteal) effect on stress-related brain activity, nor the interaction between the two factors with cluster-level FWE-correction at *p* < 0.05.

### Region-of-Interest Analysis

#### Overall effect of stress

For the extracted ROIs, activation effects for the stress task emerged in the right IFG (*t* = 4.96, *p_corr_* < 0.001), the right insula (*t* = 2.64, *p_corr_* = 0.043), and the right STG (*t* = 13.16, *p_corr_* < 0.001), and deactivation effects in the left dorsal striatum (caudate nucleus) (*t* = 4.11, *p_corr_* < 0.001) and the bilateral dlPFC (*t* = 6.52, *p_corr_* < 0.001). No activation or deactivation effects emerged in the left insula, parahippocampal gyrus, and thalamus (all *p* > 0.8). For detailed statistics including t-values and p-values for all ROIs, please refer to the Supplementary Table S4.

#### No main or interaction effect of group and phase

For the main effect of group or phase, or the interaction between the two, we did not find any significant activations or deactivations in the selected ROIs (all *p_corr_* > 0.7). Please refer to Supplementary Table S4 for additional information on t-values and p-values.

### Stress-induced brain activity and PMS symptom severity

#### Whole-brain analysis with DRSP as additional predictor

On the whole-brain level, we found no main effect for DRSP. For the interaction between group and DRSP we found one significant cluster spanning the right IFG (cluster size: 302 vox; peak coordinates x = 52, y = 22, z = 6; *t* = 4.69; cluster-level *p_FWE_* = 0.001) for the positive interaction, and one significant cluster spanning the right superior temporal gyrus (STG) and the Heschl gyrus (cluster size: 256 vox; peak coordinates x = 50, y = −16, z = 8; *t* = 4.54; cluster-level *p_FWE_* = 0.004) for the negative interaction. More specifically, we found that there was a positive association between activity in the right IFG and the mean luteal DRSP score in the control group irrespective of cycle phase, while this effect was reversed in the PMS group. For the right STG, there was a negative association between activity and mean DRSP score in the control group, and no association in the PMS group. For an illustration of the effect, see Figure 5.

**Figure 5.**
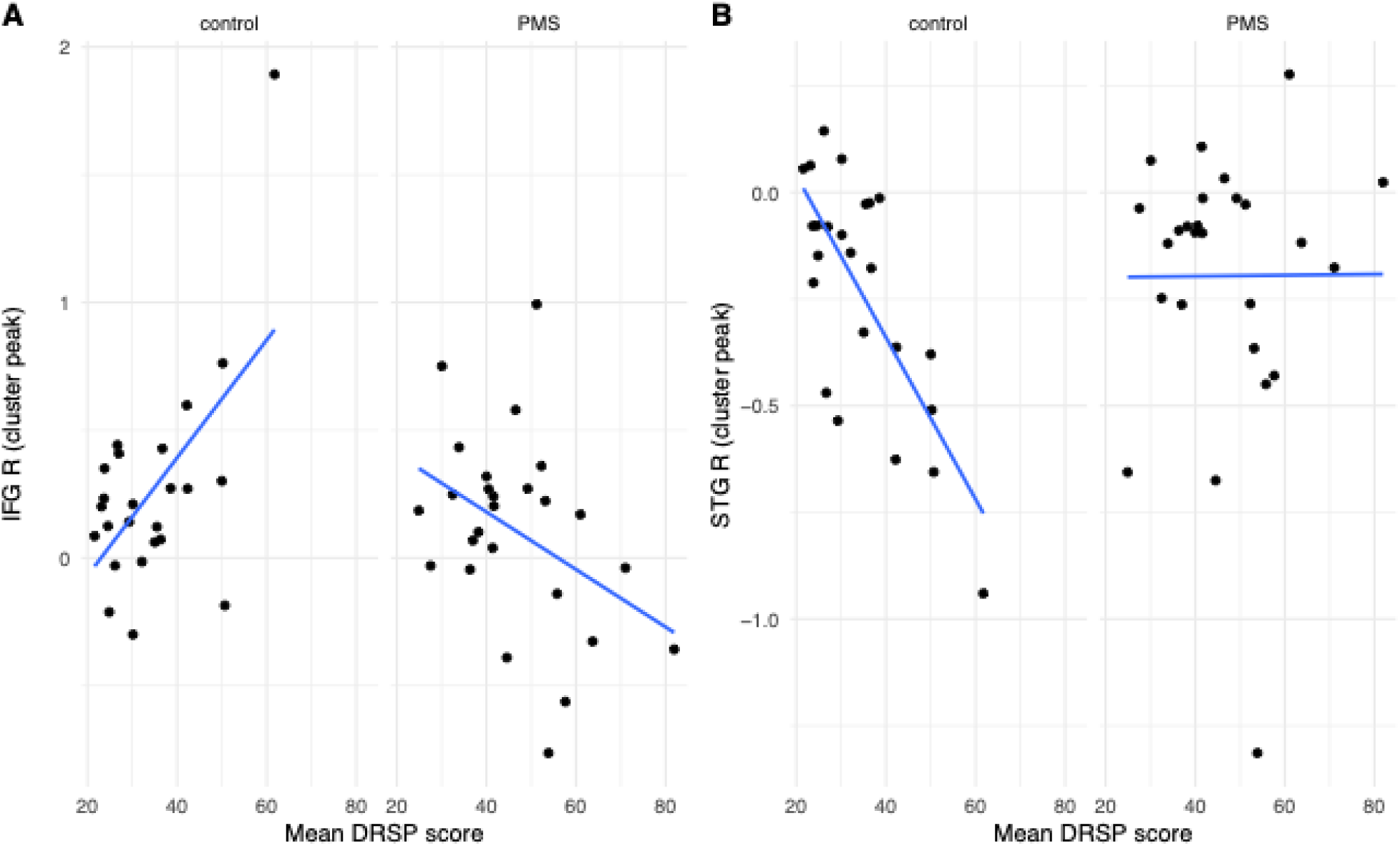
Interaction effect between group and DRSP scores (whole-brain analysis). ***Notes.*** Illustration for the interaction effect between group and DRSP scores. Figure A shows the positive interaction in the right IFG (cluster size: 302 vox; peak coordinates x = 52, y= 22, z = 6; *t* = 4.69), and Figure B shows the negative interaction in the right STG (cluster size: 256 vox; peak coordinates x = 50, y = −16, z = 8; t = 4.54). DRSP = Daily Record of Severity of Problems; IFG = inferior frontal gyrus; PMS = premenstrual syndrome; R = right; STG = superior temporal gyrus.

#### ROI analysis with DRSP as additional predictor

For the selected ROIs, the linear mixed models revealed a significant effect of DRSP (estimate = 0.03, *t* = 2.14, *p_uncorr_* = 0.037) and the interaction of group and DRSP (estimate = −0.02, *t* = −2.65, *p_uncorr_* = 0.011), both in the right IFG. None of these effects survived correcting for multiple testing. For a more detailed overview of the results see Supplementary Table S5.

## Discussion

The present study investigated whether neural, behavioral and endocrine stress responses are influenced by menstrual cycle phase and premenstrual symptom severity. To investigate neural stress responses, the MIST was used as an MRI-compatible psychosocial stress task. It is worth noting that the PMS and non-PMS group differed not only in premenstrual symptom severity but also several psychological characteristics, such as chronic stress, childhood trauma exposure, trait anxiety and history of depression, broadly consistent with previous reports of individuals with more severe premenstrual symptoms (Bencker et al., 2025). While subjective stress did not differ significantly between menstrual cycle phases or the two groups with and without PMS symptoms, subjective stress scores were significantly higher after the MIST, indicating successful stress induction. Notably, this pattern was not reflected in salivary cortisol, as averaged cortisol levels decreased from the beginning to the end of the MRI session, but no increase before and after the MIST was detected. It is not uncommon for only a subset of participants to exhibit a cortisol response to stress. One of the first studies using the MIST divided the exposed group into responders and non-responders based on their cortisol pattern, with roughly 50% in each group (Pruessner et al., 2008). Another study comparing cortisol responses to stress between women and men found that only one third of both men and women showed a cortisol increase in response to a similar MRI-compatible stress task (Kuhn et al., 2023). Both studies nevertheless reported increases in subjective stress, suggesting that successful stress induction at the subjective level is not always mirrored at the endocrine revel. Likely, the MRI experience itself already evokes stress, which could explain the decrease in cortisol from the beginning to the end of the MRI session observed in our study. While subjective and endocrine measures yielded divergent results, we did observe neural activation and deactivation effects in regions previously implicated in (psychosocial) stress. Activations in the bilateral insula, right IFG and left STG were observed in our sample and have also been reported in two meta-analyses on psychosocial stress (Berretz et al., 2021; Kogler et al., 2015b). ROI analyses further revealed deactivations in the left dlPFC and left caudate nucleus, which have likewise been implicated in previous stress research (Kogler et al., 2015b; Rosenbaum et al., 2018b). Furthermore, whole-brain results indicated no evidence of either group (PMS vs. no PMS) or menstrual cycle phase (follicular vs. luteal) effects on the neural stress response measured with our task. Importantly, these findings were obtained using a repeated-measures design in which each participant completed the stress paradigm during both the mid-follicular and late-luteal phase. This study used a repeated-measures design which increases sensitivity to menstrual cycle-related changes by minimizing between-subject variability.

Although premenstrual symptom severity showed no direct (main) effect, whole-brain analyses revealed significant interactions between group and symptom severity in the right IFG and right STG. The interaction effect of group and symptom severity within the right IFG revealed a positive association between brain activity and symptom severity in the control group, whereas this relationship was reversed in the PMS group. Specifically, higher symptom severity was associated with higher right IFG activity in the control group, but with lower right IFG activation in the PMS group. The same effect was found on the ROI-level, albeit non-significant after correction for multiple testing. One possible interpretation is that, in participants without PMS, increasing symptom severity may lead to greater recruitment of the IFG, potentially reflecting compensatory regulatory processes. In contrast, among participants with PMS, increasing symptom severity may be associated with reduced recruitment of this region, suggesting diminished regulatory capacity or a reduced ability to compensate for stress-related demands. The IFG is considered a central node in acute stress processing, particularly during psychosocial stress (Berretz et al., 2021; Kogler et al., 2015b). It has been implicated in inhibitory control (Golde et al., 2020), rumination (Int-Veen et al., 2023; Rosenbaum et al., 2018b), and is involved in coordination with limbic regions such as the insula (Berretz et al., 2021; Kogler et al., 2015b). Interestingly, previous studies have shown that high ruminators exhibit attenuated right IFG activation during social-evaluative stress, which in turn mediates post-stress rumination and negative affect (Rosenbaum et al., 2018b). Similarly, in depressed patients, reduced right IFG activation during stress appears to be more strongly linked to state rumination than to the diagnosis per se (Int-Veen et al., 2023), and influenced by childhood trauma and social anxiety (Rosenbaum et al., 2024, 2021). Our findings are broadly consistent with this literature and suggest that the degree to which the right IFG can be recruited during psychosocial stress may be a critical factor for vulnerability or resilience. Given the strong structural and functional connections of the IFG with regions such as the insula and amygdala (Berretz et al., 2021; Gold et al., 2015; Kogler et al., 2015b), future studies should also examine functional connectivity within this network. Furthermore, as some studies have proposed neuromodulatory approaches targeting the PFC, and the IFG in particular (Moses et al., 2023), future research could investigate whether similar approaches may be relevant for PMS or PMDD.

An opposite pattern was observed in the right STG. Here, we found a negative association between neural activity and premenstrual symptom score in the control group, whereas no significant association was observed in the PMS group. In the control group, greater symptom severity was associated with lower activation, or even deactivation, of the right STG. In other words, greater premenstrual symptom severity was associated with lower activation of the right STG, whereas symptom severity was not significantly associated with right STG activation in the PMS group. The superior temporal gyrus is a key brain region for processing auditory information, especially speech input (Yi et al., 2019), but has also been implicated in responses to psychosocial stress, cognitive stress regulation, and resilience factors such as self-esteem during stress processing (Kogler et al., 2017, 2015b, 2015a). The right STG is further associated with attention processing and sensitive to sex-effects and sex-steroids (Kogler et al., 2017, 2015a). Additionally, the temporal cortex expresses a high amount of aromatase, an enzyme that converts androgens into estrogens (Denley et al., 2018), and therewith seem to be linked to sex-steroid-sensitive fluctuations. As a long-term effect, stress can disrupt the connection between the STG and the hippocampus, potentially affecting memory formation and consolidation, at least in males (Shields et al., 2019). Although the present finding should be interpreted with care, it may indicate that the normative relationship between premenstrual symptom severity and STG recruitment in response to a stressor observed in controls is altered or absent in individuals with PMS as a consequence of the symptom burden, or that a plateau of activity is reached with a certain degree of symptom severity.

Although our findings did not indicate substantial differences in neural stress responses between women with and without PMS or across menstrual cycle phases, they provide important evidence regarding the stability of neural responses to stress in relation to PMS and menstrual cycle phase. Using a repeated-measures design, we found no evidence that acute psychosocial stress responses undergo systematic within-subject changes from the mid-follicular to the late-luteal phase. This suggests that the neural stress network is relatively robust to menstrual cycle-related hormonal fluctuations, at least in women with the range of premenstrual symptom severity represented in our sample. At the same time, our findings in the IFG and STG regarding interactions between group and symptom severity could suggest that symptom severity may be associated with subtle differences in the neural recruitment of regions involved in stress processing. Interestingly, thinner cortices have been found in women with PMDD compared to healthy controls in the IFG as well as the STG, possibly linking structural to functional changes (Dubol et al., 2022).

A major strength of this study is the within-subject design, allowing us to assess the same participants during both the mid-follicular and the late-luteal phase of the menstrual cycle. Compared to between-subject designs, this approach reduces interindividual variability and can detect menstrual cycle effects and the association with premenstrual symptoms much more reliable. To date, studies investigating neural stress reactivity and menstrual cycle phase have used a cross-over design (Albert et al., 2015; Chung et al., 2016). However, the repeated-measures approach comes with some limitations. The stress task was not originally designed for repeated application, even though habituation and sensitization have been tested, with results showing comparable stress reactivity across two sessions (Velozo et al., 2021). To counterbalance this, we used a randomized approach, i.e., half of the participants had their first session during the mid-follicular phase, the other half during the late-luteal phase. We also tested whether there was a significant difference in brain activity between first and second measurement and found no significant clusters.

Another limitation of the study concerns the collection of premenstrual symptoms which were not assessed during the scanning period. Although participants were rating their premenstrual symptoms prospectively for one to two menstrual cycles, it was not always possible to conduct the MRI scans during this time. Consequently, symptom scores reflected individual premenstrual symptom severity rather than symptom burden at the time of scanning. Ideally, symptom scores would be collected at the same time as the MRI measurements to align subjective symptom burden and neural reactivity, but this was not feasible due to practical constraints and participant burden. It should also be mentioned that women with and without PMS differed also in chronic stress, childhood trauma, trait anxiety and history of depression. Although these differences are characteristic of PMS populations and difficult to avoid, they may also influence neural stress processing. Future studies could examine the contributions of these factors more directly. Moreover, neural responses were assessed during two cycle phases, but the menstrual cycle is characterized by substantial hormonal fluctuations that go beyond the two time-points measured in this study. Previous cross-sectional studies have also examined the ovulatory phase, with mixed findings regarding neural stress reactivity (Albert et al., 2015; Chung et al., 2016). Future studies could consider more fine-grained longitudinal assessments across cycle phases.

To summarize, this within-subject study investigated whether neural responses to psychosocial stress vary across menstrual cycle phases and premenstrual symptom severity. Upon measuring the same participants during both the mid-follicular and late-luteal phase, we found no overall effect of menstrual cycle phase (follicular and luteal phase) or premenstrual symptom severity on stress-related whole-brain neural activation. Together with the absence of menstrual cycle effects in the ROI analyses, these findings suggest that neural responses to acute psychosocial stress remain relatively stable across the menstrual cycle and across varying levels of premenstrual symptoms. At the same time, we observed interaction effects involving symptom severity in the right IFG and right STG at the whole-brain level, with ROI analyses also pointing toward involvement of the right IFG. Although these findings require replication, they suggest that individual differences in symptom severity might be associated with differing recruitment of cognitive systems involved in stress processing. Future studies, particularly those including women with more severe symptoms or diagnosed PMDD, as well as longitudinal and connectivity-based approaches are needed to clarify the role of premenstrual symptoms in neural stress processing.

## Acknowledgements

We would like to thank Ramona Täglich who enabled data collection as a “super user” during MRI measurements, as well as Katharina Rogg, Marieke Glöckner, Luna Lenk, Sina Wenke, Antonia Stärk, Antonia Daub, Emilia Ansorge and Lena-Sophie Hölz who helped with data collection and study organization. Thank you also to Maria Ulmer and Aaron Langkamp for their help with the salivary cortisol analysis. Further, we acknowledge the MRI Core Facility of the University of Tübingen for providing access to MRI infrastructure and for their technical and personal support. We also would like to thank all the women who participated in our study for their time and effort.

## Author Contributions

EB: conceptualization, data curation, formal analysis, investigation, methodology, project administration, software, visualization, writing – original draft; ACSK: data curation, formal analysis, methodology, software, supervision, writing – review & editing; TD: supervision, writing – review & editing; ML: methodology, resources, writing – review and editing; TJ: investigation, methodology, writing – review and editing; LK: conceptualization, methodology, writing – review & editing; BD: funding acquisition, resources, writing – review & editing; AJF: funding acquisition, resources, supervision, writing – review & editing; EC: conceptualization, supervision, writing – review & editing; ACE: conceptualization, project administration, resources, supervision, writing – review & editing.

