## Supplementary Material for "Menstrual cycle phase and premenstrual symptoms in relation with the stress response: an fMRI study"

**S1.** PMDD screening questionnaire (translated from original German version)

| *Premenstrual Dysphoric Disorder* | | | |
|---|---|---|---|
| **A** | In the majority of menstrual cycles, at least 5 symptoms must be present during the week before the onset of menstruation, improve within a few days after the onset of menstruation, and become minimal or absent during the week following menstruation. | ☐ No | ☐ Yes |
| **B** | One or more of the following symptoms must be present: | ☐ No | ☐ Yes |
| B1 | **Emotional Lability**<br>Mood swings, increased sensitivity to rejection, suddenly feeling sad or tearful | ☐ No | ☐ Yes |
| B2 | **Irritability**<br>Increased irritability, anger, more frequent conflicts with others | ☐ No | ☐ Yes |
| B3 | **Depression**<br>Depressed mood, hopelessness, self-deprecating thoughts | ☐ No | ☐ Yes |
| B4 | **Anxiety**<br>Anxiety, tension, nervousness | ☐ No | ☐ Yes |

| | | | |
|---|---|---|---|
| **C** | One (or more) of the following symptoms must also be present to reach a total of 5 symptoms in combination with the symptoms listed in Criterion B above | | |
| C1 | **Reduced interest in usual activities (work, school, friends, hobbies)** | ☐ No | ☐ Yes |
| C2 | **Difficulty concentrating** | ☐ No | ☐ Yes |
| C3 | **Fatigue, rapid exhaustion, lack of energy** | ☐ No | ☐ Yes |
| C4 | **Increased appetite, overeating, cravings for certain foods** | ☐ No | ☐ Yes |
| C5 | **Insomnia or increased need for sleep** | ☐ No | ☐ Yes |
| C6 | **Feeling overwhelmed or a loss of control** | ☐ No | ☐ Yes |
| C7 | **Physical symptoms**: breast tenderness, swollen breasts, joint or muscle pain, feeling bloated, weight gain | ☐ No | ☐ Yes |

Note: Symptoms described in criteria A-C must apply to most menstrual cycles over the past year

| | | | | | |
|---|---|---|---|---|---|
| **D** | **Suffering**<br>Do the symptoms cause significant limitations in your daily life?<br>On a scale from 1 – not at all to 4 – very much, do the symptoms cause impairment with<br>(4 – very much in at least one of A-E) | ☐ No<br>☐ No | | ☐ Yes<br>☐ Yes | |
| | A. Work/School (productivity, effectiveness) | 1 | 2 | 3 | 4 |
| | B. Relationships with coworkers | 1 | 2 | 3 | 4 |
| | C. Relationships with family members | 1 | 2 | 3 | 4 |
| | D. Social activities | 1 | 2 | 3 | 4 |
| | E. Household chores | 1 | 2 | 3 | 4 |

| | | | |
|---|---|---|---|
| **E** | **Consideration of other psychiatric disorders**<br>Are your symptoms an exacerbation of symptoms of another mental disorder—such as depression or an anxiety disorder—that also occur outside the premenstrual phase? | | |
| → | The disorder is not merely an exacerbation of the symptoms of another disorder, such as major depressive disorder, panic disorder, persistent depressive disorder (dysthymia), or a personality disorder (although it may co-occur with one of these disorders). | ☐ No | ☐ Yes |

| | | | |
|---|---|---|---|
| **G** | **Consideration of Substances and Medical Conditions**<br>In your opinion, were your symptoms related to a medical condition, medication use, or the use of drugs and alcohol? | | |
| → | The symptoms are not a direct physiological consequence of a substance (substance abuse, medication, other treatment) or a general medical condition (e.g., hyperthyroidism) | ☐ Nein | ☐ Ja |

**S2.** Salivary cortisol samples analysis

Saliva samples were immediately frozen after collection and stored at −20°C until analysis. Samples were prepared according to a standardized workflow, including slow thawing at 4°C, protein precipitation and solid-phase extraction on a 96-well plate Oasis PRIME hydrophilic–lipophilic balance material. Extracts were dried under nitrogen and reconstituted in 50 µL of MeOH-H2O (30:70; v/v) in a 96-well collection plate. Chromatographic separation was performed on a Kinetex XB-C18 column (100 mm × 1.0 mm, 2.6 µm, 100 Å pore size) with a KrudKatcher ultra in-line filter on a 1290 Infinity II LC and Multisampler. For analyte detection, a QTRAP 4500 mass spectrometer (Sciex) was used in negative ionization mode. For the quantification of cortisol and cortisone in saliva samples, external calibration in artificial saliva was established, with $^{13}C_3$-labeled cortisone and $^{13}C_3$-labeled cortisol used as internal standards. The validated method according to FDA guidelines included calibration ranges from 0.55 nmol/L to 55.52 nmol/L for cortisone and cortisol.

**S3.** Preprocessing with fmriprep

Results included in this manuscript come from preprocessing performed using *fMRIPrep*24.1.1 (Esteban et al. (2019); Esteban et al. (2018); RRID:SCR_016216), which is based on *Nipype* 1.8.6 (K. Gorgolewski et al. (2011); K. J. Gorgolewski et al. (2018); RRID:SCR_002502).

**Preprocessing of $B_0$ inhomogeneity mappings**
A total of 3 fieldmaps were found available within the input BIDS structure for this particular subject. A *$B_0$* nonuniformity map (or *fieldmap*) was estimated from the phase-drift map(s) measure with two consecutive GRE (gradient-recalled echo) acquisitions. The corresponding phase-map(s) were phase-unwrapped with prelude (FSL None).

**Anatomical data preprocessing**
A total of 2 T1-weighted (T1w) images were found within the input BIDS dataset. Each T1w image was corrected for intensity non-uniformity (INU) with N4BiasFieldCorrection (Tustison et al. 2010), distributed with ANTs 2.5.3 (Avants et al. 2008, RRID:SCR_004757). The T1w-reference was then skull-stripped with a *Nipype* implementation of the antsBrainExtraction.shworkflow (from ANTs), using OASIS30ANTs as target template. Brain tissue segmentation of cerebrospinal fluid (CSF), white-matter (WM) and gray-matter (GM) was performed on the brain-extracted T1w using fast (FSL (version unknown), RRID:SCR_002823, Zhang, Brady, and Smith 2001). An anatomical T1w-reference map was computed after registration of 2 <module 'nipype.interfaces.image' from '/opt/conda/envs/fmriprep/lib/python3.11/site-packages/nipype/interfaces/image.py'> images (after INU-correction) using mri_robust_template (FreeSurfer 7.3.2, Reuter, Rosas, and Fischl 2010). An anatomical T2w-reference map was computed after registration of 2 <module 'nipype.interfaces.image' from '/opt/conda/envs/fmriprep/lib/python3.11/site-packages/nipype/interfaces/image.py'> images (after INU-correction) using mri_robust_template (FreeSurfer 7.3.2, Reuter, Rosas, and Fischl 2010). Brain surfaces were reconstructed using recon-all (FreeSurfer 7.3.2, RRID:SCR_001847, Dale, Fischl, and Sereno 1999), and the brain mask estimated previously was refined with a custom variation of the method to reconcile ANTs-derived and FreeSurfer-derived segmentations of the cortical

gray-matter of Mindboggle (RRID:SCR_002438, Klein et al. 2017). A T2-weighted image was used to improve pial surface refinement. Brain surfaces were reconstructed using recon-all (FreeSurfer 7.3.2, RRID:SCR_001847, Dale, Fischl, and Sereno 1999), and the brain mask estimated previously was refined with a custom variation of the method to reconcile ANTs-derived and FreeSurfer-derived segmentations of the cortical gray-matter of Mindboggle (RRID:SCR_002438, Klein et al. 2017). Volume-based spatial normalization to two standard spaces (MNI152NLin2009cAsym, MNI152NLin6Asym) was performed through nonlinear registration with antsRegistration (ANTs 2.5.3), using brain-extracted versions of both T1w reference and the T1w template. The following templates were were selected for spatial normalization and accessed with *TemplateFlow* (24.2.0, Ciric et al. 2022): *ICBM 152 Nonlinear Asymmetrical template version 2009c* [Fonov et al. (2009), RRID:SCR_008796; TemplateFlow ID: MNI152NLin2009cAsym], *FSL's MNI ICBM 152 non-linear 6th Generation Asymmetric Average Brain Stereotaxic Registration Model* [Evans et al. (2012), RRID:SCR_002823; TemplateFlow ID: MNI152NLin6Asym].

**Functional data preprocessing**

For each of the 6 BOLD runs found per subject (across all tasks and sessions), the following preprocessing was performed. First, a reference volume was generated, using a custom methodology of *fMRIPrep*, for use in head motion correction. Head-motion parameters with respect to the BOLD reference (transformation matrices, and six corresponding rotation and translation parameters) are estimated before any spatiotemporal filtering using mcflirt (FSL , Jenkinson et al. 2002). The estimated *fieldmap* was then aligned with rigid-registration to the target EPI (echo-planar imaging) reference run. The field coefficients were mapped on to the reference EPI using the transform. The BOLD reference was then co-registered to the T1w reference using bbregister (FreeSurfer) which implements boundary-based registration (Greve and Fischl 2009). Co-registration was configured with six degrees of freedom. The aligned T2w image was used for initial co-registration.Several confounding time-series were calculated based on the *preprocessed BOLD*: framewise displacement (FD), DVARS and three region-wise global signals. FD was computed using two formulations following Power (absolute sum of relative motions, Power et al. (2014)) and Jenkinson (relative root mean square displacement between affines, Jenkinson et al. (2002)). FD and DVARS are calculated for each functional run, both using their implementations in *Nipype* (following the definitions by Power et al. 2014). The three global signals are extracted within the CSF, the WM, and the whole-brain masks. Additionally, a set of physiological regressors were extracted to allow for component-based noise correction (*CompCor*, Behzadi et al. 2007). Principal components are estimated after high-pass filtering the *preprocessed BOLD* time-series (using a discrete cosine filter with 128s cut-off) for the two *CompCor* variants: temporal (tCompCor) and anatomical (aCompCor). tCompCor components are then calculated from the top 2% variable voxels within the brain mask. For aCompCor, three probabilistic masks (CSF, WM and combined CSF+WM) are generated in anatomical space. The implementation differs from that of Behzadi et al. in that instead of eroding the masks by 2 pixels on BOLD space, a mask of pixels that likely contain a volume fraction of GM is subtracted from the aCompCor masks. This mask is obtained by dilating a GM mask extracted from the FreeSurfer's *aseg* segmentation, and it ensures components are not extracted from voxels containing a minimal fraction of GM. Finally, these masks are resampled into BOLD space and binarized by thresholding at 0.99 (as in the original implementation). Components are also calculated separately within the WM and CSF masks. For each CompCor decomposition, the *k* components with the largest singular values are retained, such that the retained components' time series are sufficient to explain 50 percent of variance across the nuisance mask (CSF, WM, combined, or temporal). The remaining components are dropped from consideration. The head-motion estimates calculated in the correction step were also placed within the corresponding confounds file. The confound time series derived from head motion estimates and global signals were expanded with the inclusion of temporal derivatives and quadratic terms for each (Satterthwaite et al. 2013). Frames that exceeded a threshold of 0.5 mm FD or 1.5 standardized DVARS were annotated as motion outliers. Additional nuisance timeseries are calculated by means of principal components analysis of the signal found within a thin band (*crown*) of voxels around the edge of the brain, as proposed by (Patriat, Reynolds, and Birn 2017). All resamplings can be performed with *a single interpolation step* by composing all the pertinent transformations (i.e. head-motion transform matrices, susceptibility distortion correction when available, and co-registrations to anatomical and output spaces). Gridded (volumetric) resamplings were performed using nitransforms, configured with cubic B-spline interpolation.

Many internal operations of *fMRIPrep* use *Nilearn* 0.10.4 (Abraham et al. 2014, RRID:SCR_001362), mostly within the functional processing workflow. For more details of the pipeline, see the section corresponding to workflows in *fMRIPrep*'s documentation.

**Copyright Waiver**
The above boilerplate text was automatically generated by fMRIPrep with the express intention that users should copy and paste this text into their manuscripts *unchanged*. It is released under the CC0 license.

### Supplementary Tables

**Table S1.1.** Results from linear mixed model predicting subjective stress ratings

| | ***ß*** | **SE** | **df** | ***t*** | ***p*** |
|---|---|---|---|---|---|
| (Intercept) | 1.79 | 0.548 | 64.57 | 3.26 | **0.002**** |
| group | 0.49 | 0.334 | 58.92 | 1.48 | 0.145 |
| phase | -0.13 | 0.335 | 540.98 | -0.39 | 0.700 |
| factor(timepoint)2 | 0.11 | 0.181 | 540.96 | 0.62 | 0.539 |
| factor(timepoint)3 | 2.91 | 0.181 | 540.96 | 16.06 | **< 0.0001***** |
| factor(timepoint)4 | 0.18 | 0.182 | 541.02 | 0.98 | 0.326 |
| factor(timepoint)5 | -0.42 | 0.181 | 540.96 | -2.30 | **0.021*** |
| factor(timepoint)6 | -0.56 | 0.182 | 541.02 | -3.07 | **0.002**** |
| group*phase | 0.25 | 0.210 | 541.00 | 1.17 | 0.241 |

*Notes.* * < 0.1, ** < 0.001, *** < 0.0001. The model was calculated as *mean_stress ~ group * phase + factor(timepoint) + (1 | participant_id)*. Betas are therefore also on a log scale. Phase was included with follicular as baseline.

**Table S1.2.** Results from pairwise comparison regarding timepoints of stress ratings

| **Contrast** | ***ß*** | **SE** | **df** | $t_{ratio}$ | $p_{corr}$ |
|---|---|---|---|---|---|
| t1-t2 | -0.11 | 0.181 | 541 | -0.62 | 0.990 |
| t1-t3 | -2.91 | 0.181 | 541 | -16.06 | **<.0001***** |
| t1-t4 | -0.18 | 0.182 | 541 | -0.98 | 0.923 |
| t1-t5 | 0.42 | 0.181 | 541 | 2.30 | 0.194 |
| t1-t6 | 0.56 | 0.182 | 541 | 3.07 | **0.027*** |
| t2-t3 | -2.80 | 0.181 | 541 | -15.45 | **<.0001***** |
| t2-t4 | -0.07 | 0.182 | 541 | -0.37 | 0.999 |
| t2-t5 | 0.53 | 0.181 | 541 | 2.92 | **0.042*** |
| t2-t6 | 0.67 | 0.182 | 541 | 3.68 | **0.003**** |
| t3-t4 | 2.73 | 0.182 | 541 | 15.04 | **<.0001***** |
| t3-t5 | 3.33 | 0.181 | 541 | 18.37 | **<.0001***** |
| t5-t6 | 3.47 | 0.182 | 541 | 19.09 | **<.0001***** |
| t4-t5 | 0.60 | 0.182 | 541 | 3.28 | **0.014*** |
| t4-t6 | 0.74 | 0.182 | 541 | 4.04 | **0.001*** |
| t5-t6 | 0.14 | 0.182 | 541 | 0.77 | 0.972 |

*Notes*. * < 0.1, ** < 0.001, *** < 0.0001. P-values were adjusted according to Tukey.

**Table S2.1.** Results from linear mixed model predicting salivary cortisol levels (log-transformed)

| | *ß* | SE | df | *t* | $p_{corr}$ |
|---|---|---|---|---|---|
| (Intercept) | 0.34 | 0.119 | 61.70 | 2.86 | **0.006*** |
| group_orig | -0.03 | 0.073 | 55.81 | -0.45 | 0.657 |
| phaseluteal | -0.17 | 0.078 | 467.81 | -2.22 | **0.027*** |
| factor(timepoint)2 | -0.07 | 0.040 | 456.45 | -1.80 | 0.072 |
| factor(timepoint)3 | -0.10 | 0.040 | 456.27 | -2.51 | **0.013*** |
| factor(timepoint)4 | -0.21 | 0.040 | 456.43 | -5.28 | **<.0001**** |
| factor(timepoint)5 | -0.24 | 0.041 | 456.95 | -5.89 | **<.0001**** |
| factor(timepoint)6 | -0.29 | 0.041 | 456.32 | -7.09 | **<.0001**** |
| group_orig:phaseluteal | 0.05 | 0.048 | 466.15 | 1.01 | 0.315 |

*Notes.* * < 0.05, ** < 0.0001. The model was calculated as *$log_{10}$(cortisol) ~ group * phase + factor(timepoint) + (1 | participant_id)*. Betas are therefore also on a log scale. Phase was included with follicular as baseline.

**Table S2.2.** Results from pairwise comparison regarding timepoints of salivary cortisol levels

| **Contrast** | *ß* | SE | df | $t_{ratio}$ | $p_{corr}$ |
|---|---|---|---|---|---|
| t1-t2 | 0.07 | 0.040 | 457 | 1.80 | 0.464 |
| t1-t3 | 0.10 | 0.040 | 457 | 2.51 | 0.125 |
| t1-t4 | 0.21 | 0.040 | 457 | 5.28 | **<.0001***** |
| t1-t5 | 0.24 | 0.041 | 458 | 5.89 | **<.0001***** |
| t1-t6 | 0.29 | 0.041 | 457 | 7.09 | **<.0001***** |
| t2-t3 | 0.03 | 0.040 | 456 | 0.70 | 0.982 |
| t2-t4 | 0.14 | 0.040 | 457 | 3.46 | **0.008*** |
| t2-t5 | 0.17 | 0.041 | 457 | 4.10 | **0.001*** |
| t2-t6 | 0.22 | 0.041 | 458 | 5.29 | **<.0001***** |
| t3-t4 | 0.11 | 0.040 | 457 | 2.77 | 0.064 |
| t3-t5 | 0.14 | 0.041 | 458 | 3.42 | **0.009*** |
| t5-t6 | 0.19 | 0.041 | 458 | 4.62 | **0.0001**** |
| t4-t5 | 0.03 | 0.041 | 457 | 0.70 | 0.982 |
| t4-t6 | 0.08 | 0.042 | 458 | 1.90 | 0.401 |
| t5-t6 | 0.05 | 0.042 | 458 | 1.18 | 0.844 |

*Notes*. * < 0.1, ** < 0.001, * < 0.0001. P-values were adjusted according to Tukey.

**Table S3.1** Overview of whole brain voxel-wise **activation** in full-factorial model ($p_{cFWE}$<.05)

| Cluster and anatomical location | side | MNI Coordinates | | | t-value |
|---|---|---|---|---|---|
| | | X | Y | Z | |
| **Cluster 1 (58236 vox)** | | | | | |
| Fusiform gyrus | L | -24 | -80 | -12 | 16.56 |
| Superior occipital gyrus | R | 18 | -96 | 16 | 15.26 |
| Cuneus | R | 14 | -96 | 8 | 15.06 |
| Middle occipital gyrus | L | -48 | -74 | 8 | 13.99 |
| Supramarginal gyrus | R | 62 | -42 | 24 | 13.78 |
| Middle temporal gyrus | R | 50 | -64 | 8 | 13.26 |
| Calcarine gyrus | L | -8 | -90 | -8 | 12.95 |
| **Cluster 2 (29881.2 vox)** | | | | | |
| Insula | R | 32 | 20 | -14 | 7.82 |
| IFG (pars Opercularis) | R | 42 | 16 | 4 | 5.80 |
| **Cluster 3 (366 vox)** | | | | | |
| Insula | L | -32 | 18 | -14 | 7.49 |
| **Cluster 4 (277 vox)** | | | | | |
| Middle temporal gyrus | L | -56 | -8 | -10 | 4.42 |
| Superior temporal gyrus | L | -58 | 0 | -14 | 4.15 |

*Notes*. Local maxima were anatomically labeled using the Anatomy Toolbox. Only peaks assigned to a probabilistic anatomical map are reported. Where multiple peaks were assigned to the same anatomical region, only the peak with the highest t-value is listed. MNI = Montreal Neurological Institute space; IFG = inferior frontal gyrus

**Table S3.2** Overview of whole brain voxel-wise **deactivation** in full-factorial model ($p_{cFWE}$<.05)

| Cluster and anatomical location | side | MNI Coordinates | | | t-value |
|---|---|---|---|---|---|
| | | X | Y | Z | |
| **Cluster 1 (7634 vox)** | | | | | |
| IFG (pars Triangularis) | L | -52 | 34 | 20 | 9.14 |
| Middle orbital gyrus | L | -24 | 30 | -18 | 8.48 |
| Middle frontal gyrus | L | -52 | 24 | 34 | 8.48 |
| **Cluster 2 (2822 vox)** | | | | | |
| Rolandic operculum | R | 40 | -16 | 18 | 8.91 |
| Superior temporal gyrus | R | 46 | -16 | 0 | 8.17 |
| Heschl Gyrus | R | 36 | -26 | 14 | 6.11 |
| Superior orbital gyrus | R | 20 | 30 | -16 | 5.80 |
| **Cluster 3 (1852 vox)** | | | | | |
| Angular gyrus | L | -38 | -70 | 50 | 8.64 |
| Inferior parietal lobule | L | -34 | -70 | 46 | 8.56 |
| **Cluster 4 (1067 vox)** | | | | | |
| Postcentral gyrus | L | -60 | -8 | 14 | 5.84 |
| Insula | L | -36 | -18 | 16 | 4.90 |
| Heschl gyrus | L | -32 | -28 | 14 | 4.85 |
| Precentral gyrus | L | -58 | 0 | 24 | 4.53 |
| Rolandic operculum | L | -42 | -16 | 22 | 4.45 |
| Superior temporal gyrus | L | -50 | -10 | -2 | 4.26 |
| **Cluster 5 (735 vox)** | | | | | |
| Precentral gyrus | R | 48 | -14 | 48 | 5.13 |
| Postcentral gyrus | R | 62 | -4 | 32 | 5.07 |
| **Cluster 6 (412 vox)** | | | | | |
| Angular gyrus | R | 40 | -72 | 48 | 6.08 |

| | | | | | |
|---|---|---|---|---|---|
| Inferior parietal lobule | R | 40 | -50 | 40 | 5.33 |
| **Cluster 7 (396 vox)** | | | | | |
| Cerebellum (Crus 2) | R | 38 | -66 | -42 | 7.70 |
| **Cluster 8 (346 vox)** | | | | | |
| Paracentral lobule | R | 6 | -28 | 62 | 6.34 |
| Paracentral lobule | L | -6 | -28 | 66 | 4.33 |

*Notes*. Local maxima were anatomically labeled using the Anatomy Toolbox. Only peaks assigned to a probabilistic anatomical map are reported. Where multiple peaks were assigned to the same anatomical region, only the peak with the highest t-value is listed. MNI = Montreal Neurological Institute space; IFG = inferior frontal gyrus.

**Table S4.** ROI analysis

| **Contrast** | **ROI name** | **contrast value** | ***t*- value** | $p_{uncorr.}$ | $p_{corr.}$ |
|---|---|---|---|---|---|
| **Positive effect of condition = activation** | | | | | |
| | IFG R | 0.93 | 4.96 | **<0.001**** | **<0.001**** |
| | Insula L | 0.2 | 0.95 | 0.173 | 0.818 |
| | Insula R | 0.62 | 2.64 | **0.005*** | **0.043*** |
| | Parahippocampal gyrus | -0.08 | -0.31 | 0.621 | 1 |
| | STG R | 2.2 | 13.16 | **<0.001**** | **<0.001**** |
| | Thalamus R | -0.09 | -0.93 | 0.822 | 1 |
| | Caudate nucleus L | -0.69 | -4.11 | 1 | 1 |
| | dlPFC | -1.33 | -6.52 | 1 | 1 |
| **Negative effect of condition = deactivation** | | | | | |
| | IFG R | -0.93 | -4.96 | 1 | 1 |
| | Insula L | -0.2 | -0.95 | 0.827 | 1 |
| | Insula R | -0.62 | -2.64 | 0.995 | 1 |
| | Parahippocampal gyrus | 0.08 | 0.31 | 0.379 | 0.986 |
| | STG R | -2.2 | -13.16 | 1 | 1 |
| | Thalamus R | 0.09 | 0.93 | 0.178 | 0.829 |
| | Caudate nucleus L | 0.69 | 4.11 | **<0.001**** | **<0.001**** |
| | dlPFC | 1.33 | 6.52 | **<0.001**** | **<0.001**** |
| **Main effect of group** | | | | | |
| | IFG R | 0 | 0.02 | 0.885 | 1 |
| | Insula L | 0.01 | 0.04 | 0.84 | 1 |
| | Insula R | 0 | 0.01 | 0.927 | 1 |
| | Parahippocampal gyrus | 0.45 | 1.31 | 0.256 | 0.93 |
| | STG R | 0.01 | 0.05 | 0.825 | 1 |
| | Thalamus R | 0.01 | 0.29 | 0.59 | 1 |
| | Caudate nucleus L | 0 | 0 | 0.96 | 1 |
| | dlPFC | 0.02 | 0.11 | 0.742 | 1 |
| **Main effect of phase** | | | | | |
| | IFG R | 0.01 | 0.03 | 0.86 | 1 |
| | Insula L | 0.05 | 0.24 | 0.624 | 1 |
| | Insula R | 0.01 | 0.03 | 0.869 | 1 |

| | | | | | |
|---|---|---|---|---|---|
| | Parahippocampal gyrus | 0.1 | 0.29 | 0.591 | 1 |
| | STG R | 0.15 | 1.12 | 0.292 | 0.955 |
| | Thalamus R | 0.03 | 0.57 | 0.451 | 0.995 |
| | Caudate nucleus L | 0.08 | 0.56 | 0.456 | 0.996 |
| | dlPFC | 0.03 | 0.14 | 0.71 | 1 |
| **Interaction group x phase** | | | | | |
| | IFG R | 0.09 | 0.51 | 0.479 | 0.997 |
| | Insula L | 0.48 | 2.23 | 0.138 | 0.738 |
| | Insula R | 0.27 | 1.01 | 0.318 | 0.968 |
| | Parahippocampal gyrus | 0.2 | 0.59 | 0.443 | 0.995 |
| | STG R | 0.06 | 0.44 | 0.509 | 0.998 |
| | Thalamus R | 0.01 | 0.18 | 0.675 | 1 |
| | Caudate nucleus L | 0.1 | 0.7 | 0.404 | 0.991 |
| | dlPFC | 0.03 | 0.17 | 0.681 | 1 |

*Notes.* * < 0.1, ** < 0.001. Models were calculated with MarsBaR in SPM. dlPFC = dorsolateral prefrontal cortex; DRSP = Daily Rating of Severity of Problems; IFG = inferior frontal gyrus; L = left; R = right; ROI = region of interest; STG = superior temporal gyrus.

**Table S5.** ROI analysis with group, phase and DRSP as predictors

| **Predictor** | ***ß*** | **SE** | **t** | **df** | $p_{corr}$ | **ROI (outcome)** |
|---|---|---|---|---|---|---|
| (Intercept) | 0.297 | 0.195 | 1.527 | 74.865 | 0.131 | IFG R |
| group | 0.012 | 0.103 | 0.118 | 46 | 0.906 | IFG R |
| phase | -0.014 | 0.066 | -0.215 | 49 | 0.831 | IFG R |
| DRSP | 0.029 | 0.013 | 2.144 | 46 | **0.037*** | IFG R |
| group*DRSP | -0.021 | 0.008 | -2.653 | 46 | **0.011*** | IFG R |
| (Intercept) | -0.055 | 0.22 | -0.248 | 82.252 | 0.804 | Insula L |
| group | 0.063 | 0.111 | 0.565 | 46 | 0.575 | Insula L |
| phase | 0.034 | 0.083 | 0.406 | 49 | 0.687 | Insula L |
| DRSP | 0.016 | 0.014 | 1.138 | 46 | 0.261 | Insula L |
| group*DRSP | -0.013 | 0.008 | -1.584 | 46 | 0.12 | Insula L |
| (Intercept) | 0.208 | 0.249 | 0.834 | 76.522 | 0.407 | Insula R |
| group | 0.021 | 0.13 | 0.159 | 46 | 0.875 | Insula R |
| phase | -0.018 | 0.086 | -0.206 | 49 | 0.838 | Insula R |
| DRSP | 0.026 | 0.017 | 1.502 | 46 | 0.14 | Insula R |
| group*DRSP | -0.019 | 0.01 | -1.908 | 46 | 0.063 | Insula R |
| (Intercept) | 0.73 | 0.187 | 3.904 | 65.757 | **< 0.001**** | STG R |
| group | -0.027 | 0.104 | -0.261 | 46 | 0.796 | STG R |
| phase | -0.064 | 0.053 | -1.197 | 49 | 0.237 | STG R |
| DRSP | 0.022 | 0.014 | 1.611 | 46 | 0.114 | STG R |
| group*DRSP | -0.014 | 0.008 | -1.799 | 46 | 0.079 | STG R |
| (Intercept) | -0.384 | 0.177 | -2.165 | 73.923 | **0.034*** | Caudate nuc. L |
| group | 0.107 | 0.094 | 1.14 | 46 | 0.26 | Caudate nuc. L |
| phase | 0.044 | 0.059 | 0.754 | 49 | 0.455 | Caudate nuc. L |
| DRSP | 0 | 0.012 | 0.013 | 46 | 0.99 | Caudate nuc. L |
| group*DRSP | -0.006 | 0.007 | -0.875 | 46 | 0.386 | Caudate nuc. L |

| | | | | | | |
|---|---|---|---|---|---|---|
| (Intercept) | -0.203 | 0.229 | -0.885 | 67.635 | 0.379 | dlPFC |
| group | -0.05 | 0.126 | -0.399 | 46 | 0.692 | dlPFC |
| phase | -0.027 | 0.068 | -0.393 | 49 | 0.696 | dlPFC |
| DRSP | 0.008 | 0.016 | 0.464 | 46 | 0.645 | dlPFC |
| group*DRSP | -0.004 | 0.01 | -0.44 | 46 | 0.662 | dlPFC |
| (Intercept) | -0.063 | 0.101 | -0.617 | 87.76 | 0.539 | Thalamus R |
| group | 0.001 | 0.049 | 0.022 | 46 | 0.983 | Thalamus R |
| phase | 0.027 | 0.041 | 0.666 | 49 | 0.509 | Thalamus R |
| DRSP | 0 | 0.006 | -0.054 | 46 | 0.957 | Thalamus R |
| group*DRSP | -0.001 | 0.004 | -0.355 | 46 | 0.724 | Thalamus R |
| (Intercept) | 0.214 | 0.285 | 0.75 | 78.339 | 0.455 | Parahipp. |
| group | -0.099 | 0.148 | -0.668 | 46 | 0.507 | Parahipp. |
| phase | -0.056 | 0.102 | -0.549 | 49 | 0.586 | Parahipp. |
| DRSP | -0.002 | 0.019 | -0.128 | 46 | 0.898 | Parahipp. |
| group*DRSP | -0.001 | 0.011 | -0.112 | 46 | 0.912 | Parahipp. |

*Notes.* * < 0.1, ** < 0.001. Linear mixed models were calculated as *ROI ~ group + phase + DRSP + DRSP*group + (1 | subject)*. dlPFC = dorsolateral prefrontal cortex; DRSP = Daily Rating of Severity of Problems; IFG = inferior frontal gyrus; L = left; nuc. = nucleus; Parahipp. = parahippocampal gyrus; R = right; ROI = region of interest; STG = superior temporal gyrus.
